# A GWAS–machine learning framework reveals protein-synthesis pathway signals for yield in *Theobroma cacao* after population-structure correction

**DOI:** 10.1101/2025.04.23.650203

**Authors:** Insuck Baek, Jishnu Bhatt, Seunghyun Lim, Dongho Lee, Jae Hee Jang, Stephen P. Cohen, Amelia H. Lovelace, Moon S. Kim, Lyndel W. Meinhardt, Sunchung Park, Ezekiel Ahn

## Abstract

Improving cacao yield, a key objective in post-domestication crop improvement, remains a primary goal for breeders, but progress is often hindered by the confounding effects of population structure. To overcome this, we analyzed 346 diverse cacao accessions using an ML-based association mapping framework (with and without population structure adjustment) and a phenotype-only ML prediction of yield. By correcting for population structure, our Bootstrap Forest-based GWAS revealed association signals that showed consistent enrichment for ribosome and protein-synthesis functions, and a recurrent subset of SNPs with high importance appeared across multiple yield components, including pod index and seed number. In parallel, a Neural Network model was utilized to identify cotyledon mass and length as the most powerful predictors for total wet bean mass (R² = 0.715 by repeated five-fold cross-validation), suggesting a practical, low-cost screening proxy for breeding). Collectively, this study delivers a robust genetic framework and a novel predictive tool to accelerate the development of high-yielding cacao varieties through the early identification of elite clones.

## Introduction

Improving the genetic potential of high-value horticultural crops is a central goal of modern agricultural science. *Theobroma cacao* L., the source of cocoa beans, is a globally significant crop that underpins a multi-billion-dollar chocolate industry and supports the livelihoods of millions of smallholder farmers^1,2^.These smallholders are responsible for approximately 80% of global cocoa production^3^. Originating in the upper Amazon basin^4^, cacao exhibits rich genetic diversity, contributing to a wide range of flavors, aromas, and other desirable traits^5^. However, cacao production faces increasing challenges from emerging diseases and pests, and a growing global demand for chocolate^6,7^. These challenges necessitate the development of improved cacao varieties with enhanced yield potential, disease resistance, and environmental resilience, while maintaining and enhancing quality attributes. This continuous drive for crop improvement, from early domestication to modern breeding, requires a deep understanding of the genetic basis of key agronomic traits.

Efforts to understand genetic diversity and identify genes controlling important traits from the cacao genome have been ongoing for decades. Studies have indicated that genetic differentiation within cacao lineages results from spatial variation in selection pressures, genetic drift, or a combination of both processes^8^. Yield stability over time is a critical trait in cacao breeding^3^, yet studies focusing on temporal stability remain limited^9^. A study evaluating 34 cacao hybrid families over six years in Costa Rica revealed substantial variability in annual yield, with coefficient of variation values ranging from 113% to 642% for different years, particularly notable in years two and three, when 80% of trees produced no pods^9^. Cacao yield was contingent on interrelated factors such as pod health and disease incidence, shown through a positive correlation between the number of healthy pods and overall yield efficiency^3^. When low yield variability is optimized for in breeding programs, newly developed cultivars are susceptible to reduced yield efficiencies, necessitating broadly informed selection of genetic traits to sustain productivity over time^3^. Genetic factors significantly influence pod development and quality traits^10^. The selection of progeny based on pod index, calculated as the number of pods required to produce one kilogram of dried cocoa^11^, and weight can improve overall tree productivity and efficiency in production processes^10^. Monitoring genetic identity is essential as progenies from mislabeled or pollen-contaminated clones confound breeding efforts towards the development of superior varieties^12^.

Traditional breeding methods, relying on phenotypic selection and controlled crosses, have contributed to significant improvements in cacao varieties^13,14^. However, these methods are unable to take into account the complex interplay and inheritance of many traits^15^. The advent of genomic tools, such as genome-wide association studies (GWAS), offers new opportunities to understand inherited trait complexity and accelerate cacao improvement^16,17^. Previous studies in cacao have employed GWAS to identify loci associated with disease resistance, yield components, and bean quality attributes^11,16,17^. Despite the use of standard statistical corrections, a critical limitation of these studies is that the strong population structure in cacao often results in associations with general ‘vigor’ rather than specific, actionable genes. Furthermore, translating these genetic findings into practical tools for breeders remains a significant hurdle. Consequently, a comprehensive understanding of the genetic architecture of complex traits in cacao, such as those related to yield potential, has remained elusive. Only a few studies have integrated advanced machine learning techniques to enhance the prediction and understanding of complex traits in cacao, primarily using inputs such as image analysis for bean quality and canopy architecture^18,19^, sensor data for fermentation level^20^, or agronomic and climate data for yield potential^21^. However, studies that directly combine machine learning frameworks with high-throughput genomic data for trait discovery and prediction remain rare.

This study addresses these limitations by leveraging a large and diverse collection of 346 cacao accessions, utilizing publicly available phenotypic and genotypic data from the ICGT Cacao Germplasm Database^11^. These accessions, conserved *ex-situ* at the International Cocoa Genebank, Trinidad (ICGT), represent a broad range of genetic and phenotypic variation^11^. We employed a multifaceted approach, combining detailed phenotypic evaluations of 27 traits, encompassing flower, fruit, and seed characteristics, with genome-wide SNP genotyping, phylogenetic analysis, and advanced statistical methods. Specifically, as a key innovation of this study, we utilized a dual-model approach, performing the GWAS both with and without correction for population structure, to robustly identify significant marker-trait associations and explored the relationships among traits using hierarchical clustering^22^. Furthermore, based on various phenotypic data, we trained a machine-learning model to predict wet bean mass, a direct indicator of yield potential.

The primary objectives of this study were to: (1) investigate the relationship between genetic relatedness and phenotypic variation in a diverse collection of cacao accessions; (2) identify genomic regions and potential candidate genes associated with key traits using a Bootstrap Forest-based GWAS approach; (3) explore the relationships among traits through hierarchical clustering; and (4) develop and evaluate a machine learning model for predicting wet bean mass. We tested several key hypotheses: first, that phenotypic variation in this diverse collection would be complex and only partially explained by broad phylogenetic relationships; second, that our GWAS and machine learning models would successfully identify novel candidate genes and accurately predict yield potential, respectively; and finally, that a corrected GWAS model would provide a more robust approach for disentangling broad ancestral signals from loci with direct, trait-specific functions, which would in turn resolve into core biological pathways upon Gene Ontology (GO) analysis.

## Results

### Correlation between patristic distances and phenotypic traits

To assess whether more distantly related accessions differ more in phenotype, we computed Mantel correlations between patristic distances (from the phylogeny) and phenotypic distance matrices across 344 traits out of 346 accessions with both genotypes and phenotypes (intersection n= 342). Correlations were generally small (median *r* ≈ 0.10). After false discovery rate (FDR) correction (q< 0.05), 12 traits were significant, led by pod index (*r*= 0.25, q= 0.0019) and cotyledon width (*r*= 0.20, q= 0.0019) (Fig. S1). The Mantel correlations were generally small (median *r*= ∼0.10). The remaining traits were not significant after correction

### Visualization of phenotypic and genotypic diversity using PCA

Principal Component Analysis (PCA) was used to visualize and compare the patterns of variation within the phenotypic and genotypic datasets (Fig. 1). The PCA plot based on the phenotypic data reveals a single, diffuse cloud of points, indicating that the measured traits vary continuously across the cacao collection (Fig. 1a). The first two principal components explained 14.8% and 10.2% of the total phenotypic variation, respectively. This lack of distinct clustering suggests that no clear-cut “types” of trees can be delineated based on their physical traits alone. In contrast, the PCA based on the genotypic SNP data shows clear evidence of population structure (Fig. 1b). The accessions are organized into several distinct clusters, demonstrating that the collection is composed of different underlying ancestral groups. The first two principal components for the genotype data explained 16% and 11.8% of the total genetic variation. This visual evidence of genetic structuring compared to the continuous phenotypic variation powerfully justifies the necessity of correcting for population structure in the subsequent GWAS to avoid spurious findings.

**Fig. 1.**
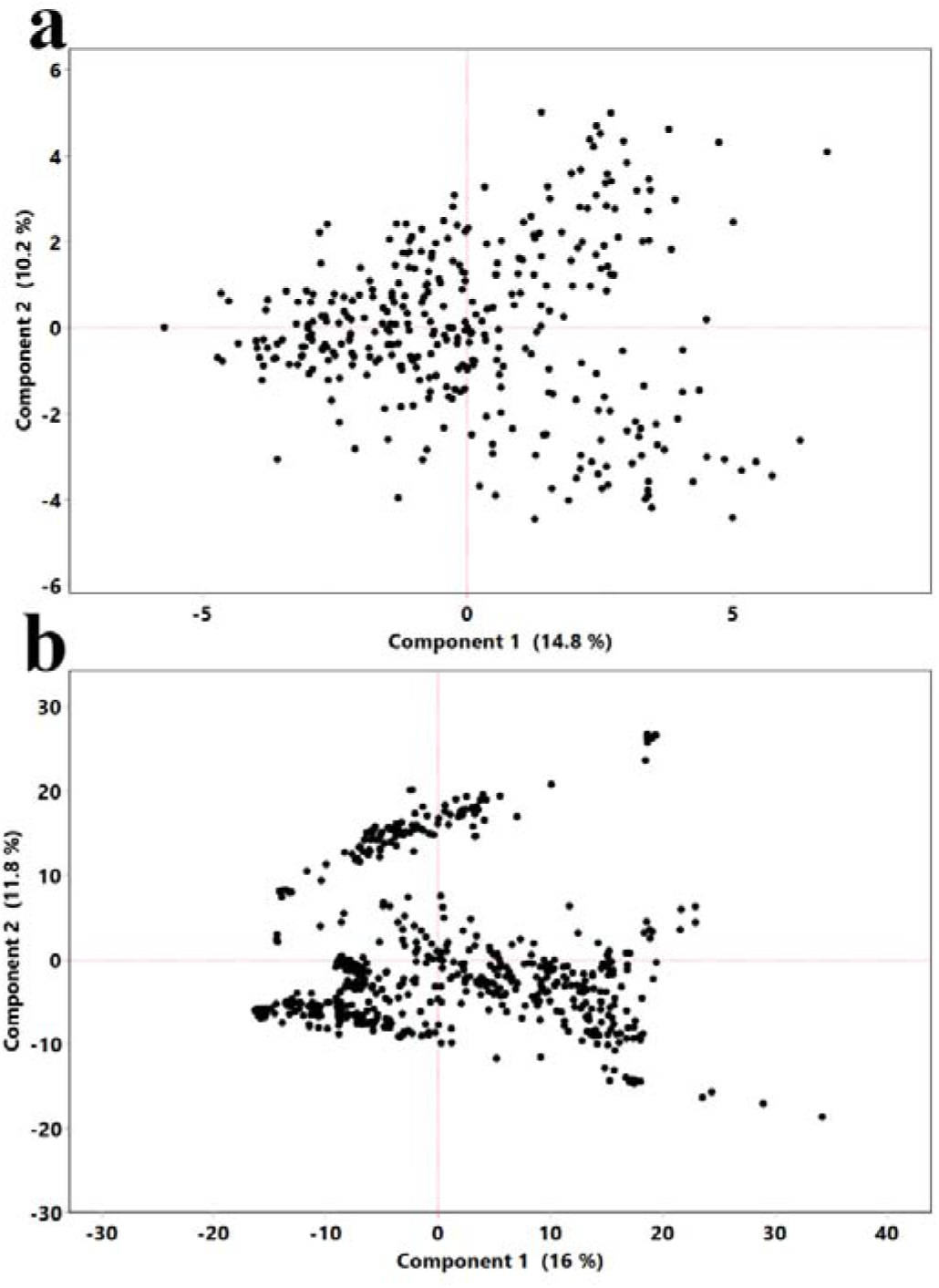
Principal component analysis of phenotypic and genotypic diversity in the cacao collection. Each point represents a single cacao accession. (a) PCA based on all measured phenotypic traits. The axes show the first two principal components, which account for 14.8% and 10.2% of the phenotypic variance, respectively. The plot reveals a continuous distribution with no distinct clusters. (b) PCA based on SNP genotype data. The axes show the first two principal components, which account for 16% and 11.8% of the genetic variance, respectively. The plot reveals clear clustering, indicating significant population structure within the collection.

### Genome-wide association study of key yield and morphological traits

To identify genomic regions associated with phenotypic variation, we employed a machine learning-based GWAS approach using Bootstrap Forest models. To specifically assess and control for the impact of the strong population structure within the collection, two parallel analyses were conducted: a naive model and a structured association model. The naive model utilized the raw phenotypic data to provide an exploratory view of all potential genetic associations, including those potentially confounded with ancestry. In contrast, the structured association model was performed on ancestry-corrected residuals to identify robust loci with a direct influence on traits, independent of the accessions’ genetic background. This dual approach allows for a direct comparison to distinguish robust, trait-specific associations from broader effects linked to the collection’s underlying genetic structure. The findings presented below are from the structured association model, which controls for the effects of population structure to identify direct marker-trait associations. This is a critical step in identifying the genetic targets of past and future selection for crop improvement.

The analysis revealed numerous significant SNP-trait associations. The top-ranking SNPs for seven key traits related to yield, morphology, and quality are summarized in Table 1 and Fig. 2. A list of the most significant associations (Portion > 0.005) for all 27 traits is available in Supplementary Data 1.

**Fig. 2.**
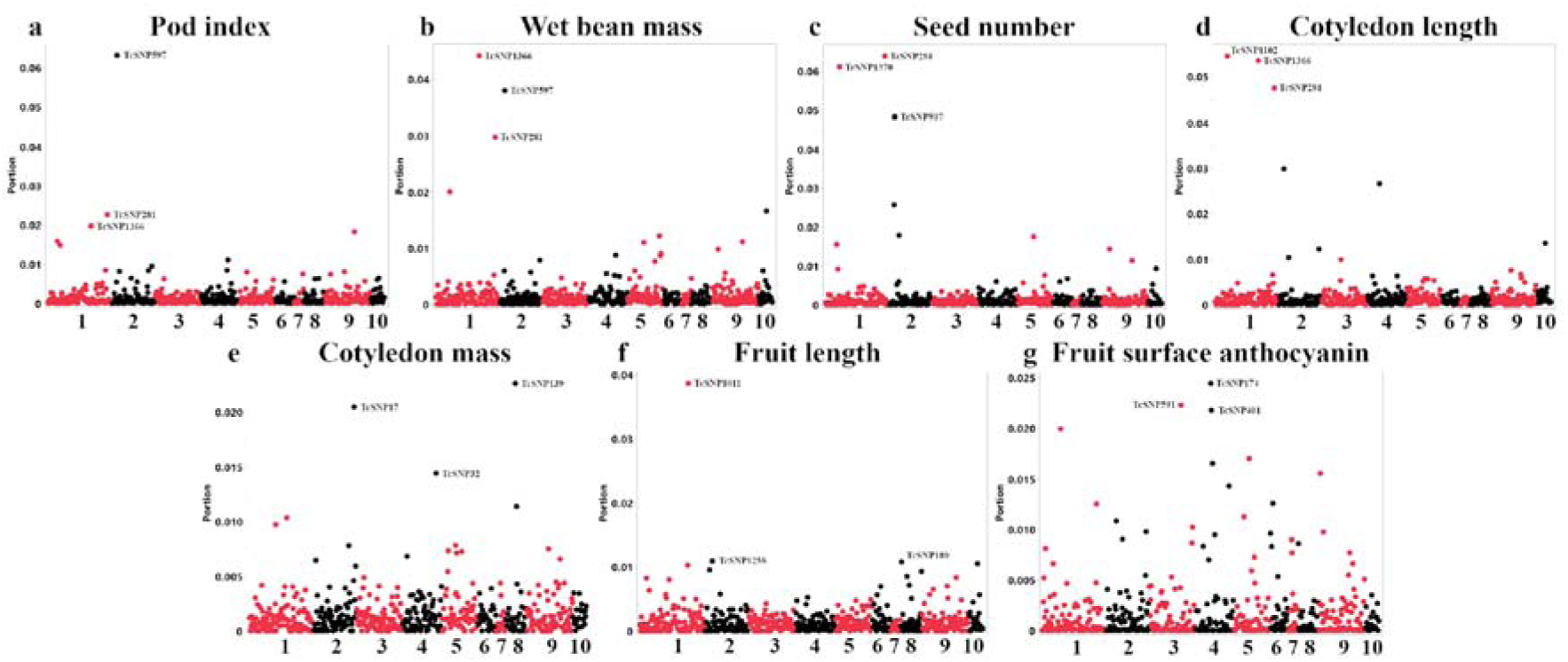
Manhattan plots from the corrected GWAS for key cacao traits. Each panel displays the results from the ancestry-adjusted Bootstrap Forest GWAS for a specific key trait: (a) Pod index, (b) Wet bean mass, (c) Seed number, (d) Cotyledon length, (e) Cotyledon mass, (f) Fruit length, and (g) Fruit surface anthocyanin. The x-axis shows the SNP position across the 10 cacao chromosomes, while the y-axis represents the importance score (Portion) from the model, where higher values indicate a stronger association. Top-ranking SNPs, corresponding to those in Table 1, are labeled with their respective IDs. SNPs with unmapped chromosomal locations are not shown.

**Table 1.** Putative candidate genes near top SNPs associated with key cacao traits from corrected GWAS. Summary of the most significant SNP associations identified through the Bootstrap Forest-based GWAS after correcting for population structure. The table lists the top-ranking SNPs for seven key traits related to yield, fruit morphology, and color. For each SNP, the table provides its chromosomal location (Chr), the nearest annotated putative gene, and the importance score (Portion) from the model. The importance score reflects the contribution of each SNP to the prediction of the trait in the Bootstrap Forest model. SNPs that could not be mapped to the reference genome are excluded.

| Trait | SNP ID | Chr | Putative Gene | Importance Score (Portion) |
| --- | --- | --- | --- | --- |
| Pod index | TcSNP597 | 2 | *Tc02v2_p004390*<br>21 kDa seed protein | 0.0632 |
|  | TcSNP281 | 1 | *Tc01v2_p034810*<br>60S ribosomal protein L31 | 0.0226 |
|  | TcSNP1366 | 1 | *Tc01v2_p025790*<br>Heparan-alpha-glucosaminide N-acetyltransferase | 0.0197 |
| Wet bean mass | TcSNP1366 | 1 | *Tc01v2_p025790*<br>Heparan-alpha-glucosaminide N-acetyltransferase | 0.0442 |
|  | TcSNP597 | 2 | *Tc02v2_p004390*<br>21 kDa seed protein | 0.038 |
|  | TcSNP281 | 1 | *Tc01v2_p034810*<br>60S ribosomal protein L31 | 0.0297 |
| Seed number | TcSNP281 | 1 | *Tc01v2_p034810*<br>60S ribosomal protein L31 | 0.0638 |
|  | TcSNP1370 | 1 | *Tc01v2_p008280*<br>Histone H3.3 | 0.0611 |
|  | TcSNP597 | 2 | *Tc02v2_p004390*<br>21 kDa seed protein | 0.0483 |
| Cotyledon length | TcSNP1102 | 1 | *Tc01v2_p006960*<br>Cytochrome c oxidase subunit 6a, mitochondrial | 0.0545 |
|  | TcSNP1366 | 1 | *Tc01v2_p025790*<br>Heparan-alpha-glucosaminide N-acetyltransferase | 0.0535 |
|  | TcSNP281 | 1 | *Tc01v2_p034810*<br>60S ribosomal protein L31 | 0.0475 |
| Cotyledon mass | TcSNP139 | 8 | *Tc08v2_p007790*<br>Putative Cytochrome B5 isoform D | 0.0221 |
|  | TcSNP17 | 2 | *Tc02v2_p029690*<br>RING/U-box superfamily protein, putative | 0.0204 |
|  | TcSNP32 | 4 | *Tc04v2_p022290*<br>Pyruvate, phosphate dikinase, chloroplastic | 0.0144 |
| Fruit length | TcSNP1011 | 1 | *Tc01v2_p028610*<br>Aspartyl protease family protein 2 | 0.0386 |
|  | TcSNP1258 | 2 | *Tc02v2_p004580*<br>Probable aquaporin PIP2-8 | 0.0109 |
|  | TcSNP189 | 8 | *Tc08v2_p000820*<br>Succinate dehydrogenase subunit 5, mitochondrial | 0.0108 |
| Fruit surface anthocyanin | TcSNP174 | 4 | *Tc04v2_p009180*<br>6-phosphogluconate dehydrogenase, decarboxylating 3 | 0.0244 |
|  | TcSNP401 | 4 | *Tc04v2_p010060*<br>ATP-dependent Clp protease proteolytic subunit 3, chloroplastic | 0.0218 |
|  | TcSNP591 | 1 | *Tc01v2_p009870*<br>60S ribosomal protein L4-1 | 0.0199 |

### Genetic loci for yield and its core components

Key yield traits, including pod index, wet bean mass, and seed number, shared a common set of associated genetic loci, highlighting a core genetic architecture for productivity. Notably, the SNPs TcSNP597 (encoding a 21 kDa seed protein) and TcSNP281 (encoding a 60S ribosomal protein L31) appeared as top associations for all three of the aforementioned key yield traits. Furthermore, TcSNP1366 (encoding a Heparan-alpha-glucosaminide N-acetyltransferase) was a top locus for both pod index and wet bean mass. These findings strongly link cacao yield potential to the genetic machinery governing protein synthesis (ribosomal proteins), energy metabolism (heparan-alpha-glucosaminide N-acetyltransferase), and the production of essential seed storage proteins.

### Associations for key predictors of yield and fruit morphology

To identify direct targets for crop improvement, GWAS also identified loci controlling important fruit and seed characteristics that contribute to overall yield potential, such as cotyledon and fruit size. For cotyledon mass, top SNPs included TcSNP139 (within the gene for a Putative Cytochrome B5 isoform D) and TcSNP17 (within a gene for a RING/U-box superfamily protein). For cotyledon length, the most significant association was TcSNP1102, within the gene for a mitochondrial cytochrome c oxidase subunit. Together, these results suggest that the size and mass of cotyledons are influenced by genes involved in cellular respiration, energy metabolism, and protein regulation. Additionally, for fruit length, a key component of pod size, the top SNP was TcSNP1011, located within a gene encoding an aspartyl protease family protein. Plant proteases are known to be involved in tissue remodeling, making this a strong candidate gene for influencing fruit development.

### Internal validation via pigmentation

For fruit surface anthocyanin level, the most significant SNP was TcSNP174. This SNP is located within the gene encoding 6-phosphogluconate dehydrogenase. This enzyme is a key component of the pentose phosphate pathway, which is directly responsible for producing the precursors required for flavonoid and anthocyanin biosynthesis, providing a direct link between an association and a known biochemical pathway controlling pigmentation in cacao.

### Genomic hotspots of trait association

To visualize the global distribution of the most significant marker-trait associations and identify genomic regions potentially under selection for multiple traits, the top three SNPs for each of the 27 traits were plotted according to their physical position on the cacao chromosomes (Fig. S2). This approach revealed several genomic regions, or “hotspots,” that contained significant associations for multiple, diverse traits. A notable pattern emerged where these hotspots were often concentrated near the boundaries of the chromosomes. This was particularly evident on chromosomes 1, 2, 5, 9, and 10 which all showed significant clusters of associations near their starting positions, while chromosomes 1, 2, 4, and 9 also showed clusters near their ends.

### Hierarchical clustering reveals functional trait groups

To understand how selection might act on suites of related traits, we conducted hierarchical clustering on the ancestry-adjusted SNP importance scores derived from the Bootstrap Forest GWAS. The resulting dendrogram visualizes the relationships between traits based on their shared genetic architecture (Fig. 3).

**Fig. 3.**
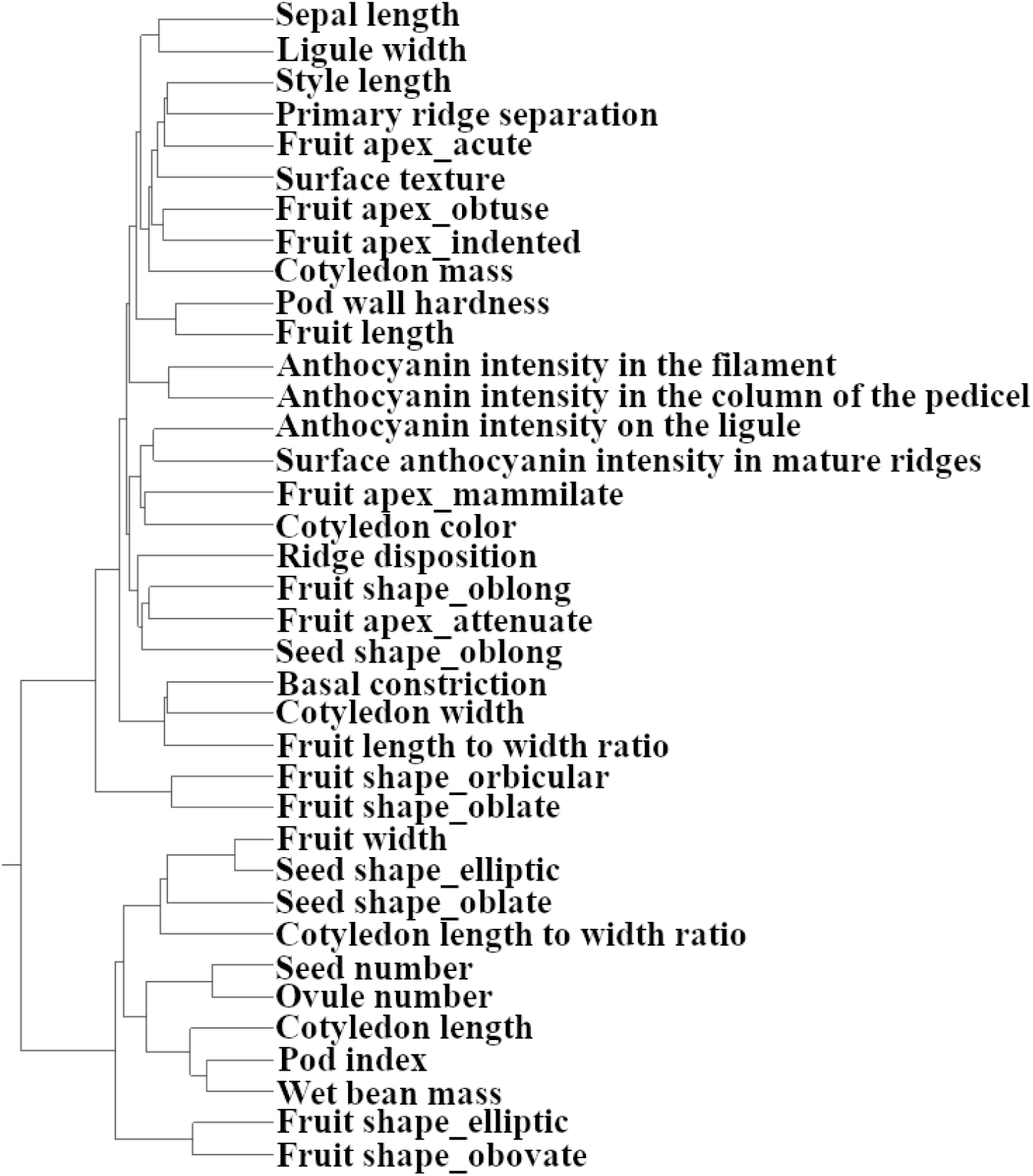
Hierarchical clustering of cacao traits based on corrected genetic association profiles. The dendrogram illustrates the relationships among traits, generated using Ward’s method on the SNP importance scores (Portion) from the corrected Bootstrap Forest GWAS. This analysis groups traits that share similar underlying genetic associations, revealing distinct functional clusters such as those related to overall yield and pigmentation.

A number of expected, functionally related traits were successfully grouped. For instance, the four measured anthocyanin intensity traits (in the filament, pedicel, ligule, and mature fruit ridges) were found in close proximity within the same parent cluster. Similarly, various fruit and seed shape traits formed logical, though not always adjacent, groupings.

The analysis revealed a distinct cluster composed of key yield-related traits. Key productivity metrics such as pod index and wet bean mass formed a tight pair, which in turn clustered with another group containing seed number, ovule number, and unexpectedly cotyledon length. Cotyledon width grouped with fruit characteristics and cotyledon length-to-width ratio grouped with seed shape traits and fruit width. Cotyledon mass was placed on a distant branch of the dendrogram, positioned in proximity to traits including pod wall hardness, fruit length, and several fruit apex form traits. The grouping of these distinct but related traits indicates a shared genetic network may govern overall yield potential in cacao.

### Comparison of gene ontology enrichment between naive and structured models

To understand the biological functions of the genes associated with key traits, we performed a GO enrichment analysis. We used pod index as a case study to demonstrate the importance of correcting for population structure in a GWAS.

The GO analysis of the top candidate genes from the naive (uncorrected) model revealed a broad mix of plausible biological pathways (Fig. 4a). The results were enriched for terms related to general energy metabolism (e.g., “respiratory chain complex,” “photosynthesis light harvesting”), protein management (e.g., “protein processing in endoplasmic reticulum”), and stress response (e.g., “response to hydrogen peroxide,” “response to heat”). This represents a classic “vigor” signal, likely reflecting the broad, fundamental differences in overall health and metabolism between the different ancestral groups in the cacao collection. Detailed KEGG pathway diagrams for key findings from this uncorrected analysis, such as protein processing in the endoplasmic reticulum and photosynthesis, are shown in Supplementary Figs. S3 and S4, respectively.

**Fig. 4.**
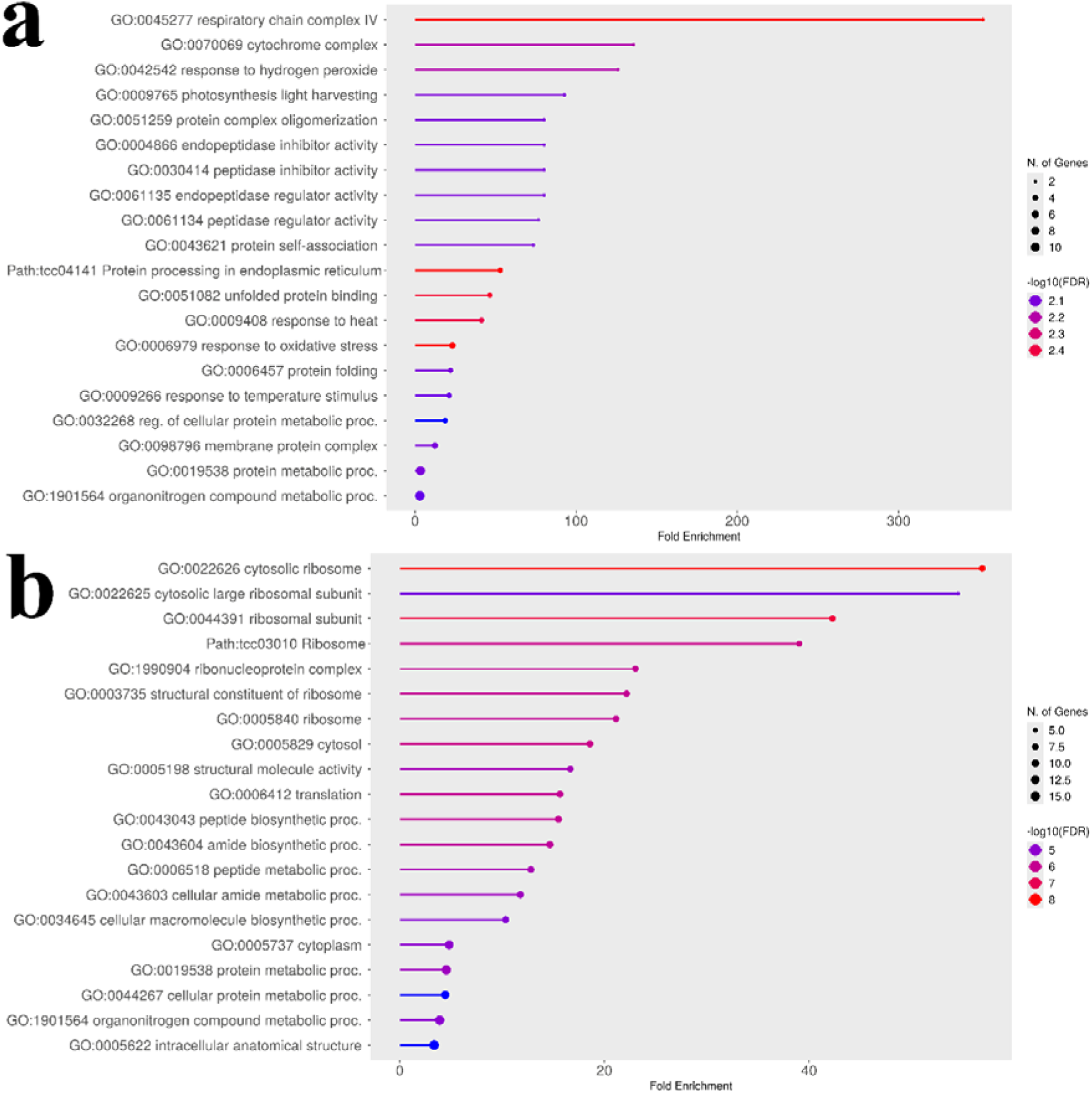
Comparison of go enrichment analyses for uncorrected and corrected pod index GWAS results. The plots show significantly enriched GO terms for the top candidate genes identified for the pod index trait. (a) GO enrichment plot for genes from the naive (uncorrected) analysis. The analysis reveals a broad mix of pathways related to energy metabolism, protein processing, and stress response. Detailed KEGG pathways for protein processing in the endoplasmic reticulum and photosynthesis are shown in Figs. S2 and S4, respectively. (b) GO enrichment plot for genes from the structured association (corrected) analysis. After correcting for population structure, the result is a significant and specific enrichment for the ribosome and protein synthesis pathways. A detailed KEGG pathway for the ribosome is shown in Fig. S5. This comparison illustrates how the statistical correction filtered out confounding signals to reveal the core machinery of protein synthesis as the primary biological process associated with pod index.

In contrast, the GO analysis of the top genes from the structured association (corrected) model tells a much more specific and statistically significant story (Fig. 4b). After correcting for population structure, the broad metabolic and stress signals disappeared, and the result became overwhelmingly dominated by a single theme: the ribosome (FDR = 1.0E-08). Multiple enriched terms, such as “cytosolic ribosome,” “ribosomal subunit,” and “translation,” all point to the core machinery of protein synthesis. A detailed KEGG pathway diagram showing the specific ribosomal components identified is provided in Fig. S5. This direct comparison demonstrates the power of the statistical correction, moving the analysis from a broad signal to a specific and robust biological insight that links pod index to the plant’s capacity for protein production.

### Combined-trait analysis reveals core biological processes

To identify the fundamental pathways underpinning groups of related traits, we performed combined GO enrichment analyses on our corrected GWAS results. First, identify the fundamental biological pathways underpinning this group of traits, we combined the candidate genes from nine size- and yield-related traits (ovule number, fruit length, fruit width, seed number, wet bean mass, cotyledon length, cotyledon width, cotyledon mass, and pod index). The GO enrichment analysis revealed a highly significant and specific enrichment for the ribosome and protein synthesis (Fig. 5a). This result indicates that the primary genetic drivers of yield are linked to the plant’s capacity for protein production.

**Fig. 5.**
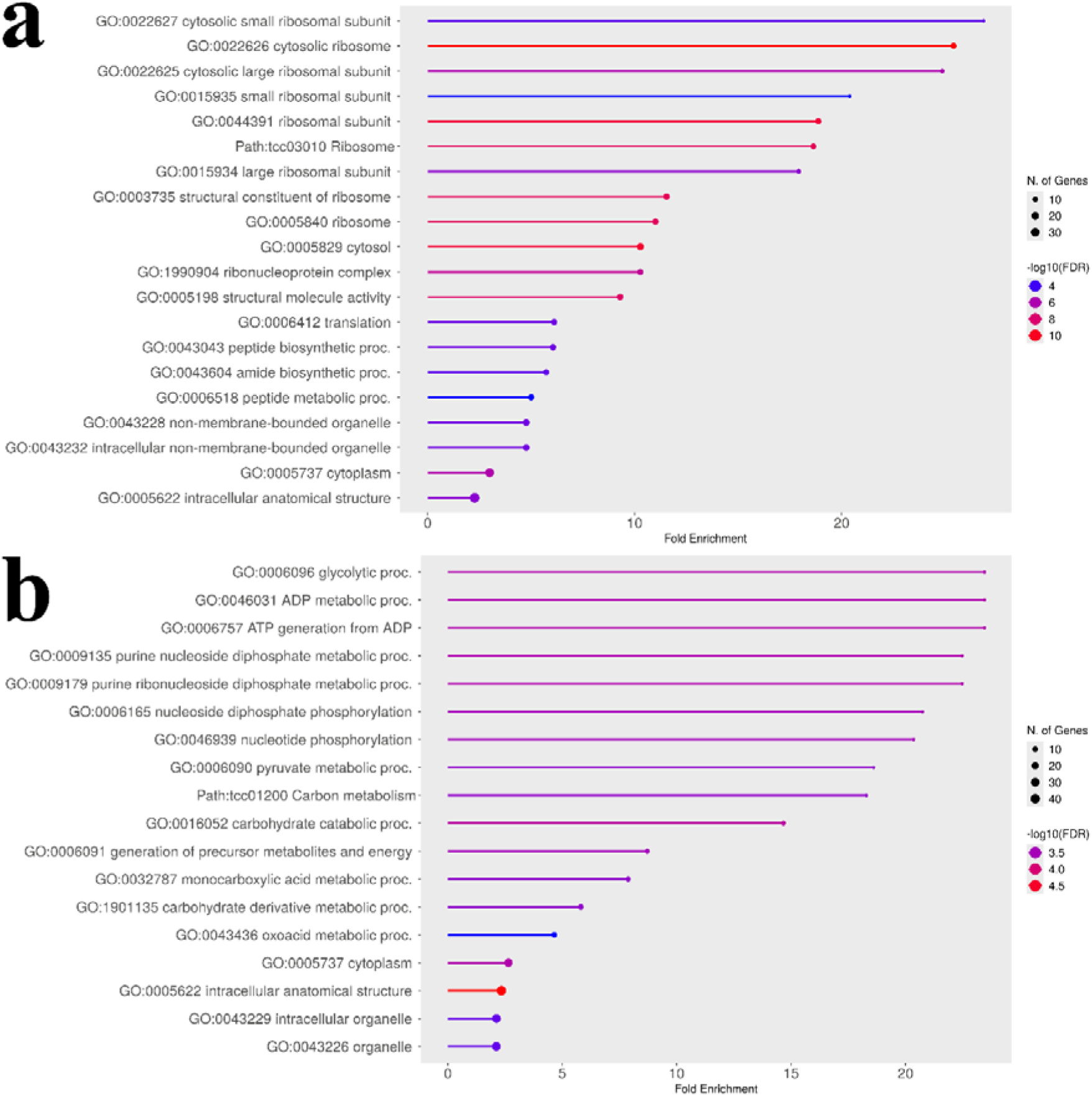
Comparison of GO enrichment plots for combined trait groups. (a) GO enrichment plot for the combined “yield & size” traits. The analysis is overwhelmingly dominated by terms related to the ribosome and protein synthesis. (b) GO enrichment plot for the combined “color & pigmentation” traits. This analysis reveals a strong enrichment for pathways related to primary energy metabolism.

To further validate our analytical process, we investigated the genetic basis of pigmentation across tissues. We performed a combined analysis of five color-related traits (flower anthocyanin intensity in the pedicel, ligule, and filament; fruit surface anthocyanin intensity; and seed cotyledon color). This analysis revealed a strong enrichment for pathways related to primary energy metabolism, including glycolysis and ATP generation (Fig. 5b), indicating that producing the anthocyanin pigments is an energetically expensive process. A key KEGG pathway identified in this analysis was “photosynthesis - antenna proteins,” and a detailed diagram showing the specific components identified, such as *Lhcb4*, is provided in Fig. S6.

A deeper look at the top KEGG pathway results provided a more nuanced story (Fig. 6). For the “yield & size” group, numerous components of both the small and large ribosomal subunits were identified (Fig. 6a). This suggests that overall yield is linked to the general capacity and throughput of the entire protein factory. In contrast, for the “color & pigmentation” group, only a few, specific ribosomal proteins were identified, such as L4e, L7e, and L24e (Fig. 6b). This suggests that the genetic control of pigmentation may be linked to these specific ribosomal proteins having specialized regulatory roles, rather than affecting the overall rate of protein synthesis.

**Fig. 6.**
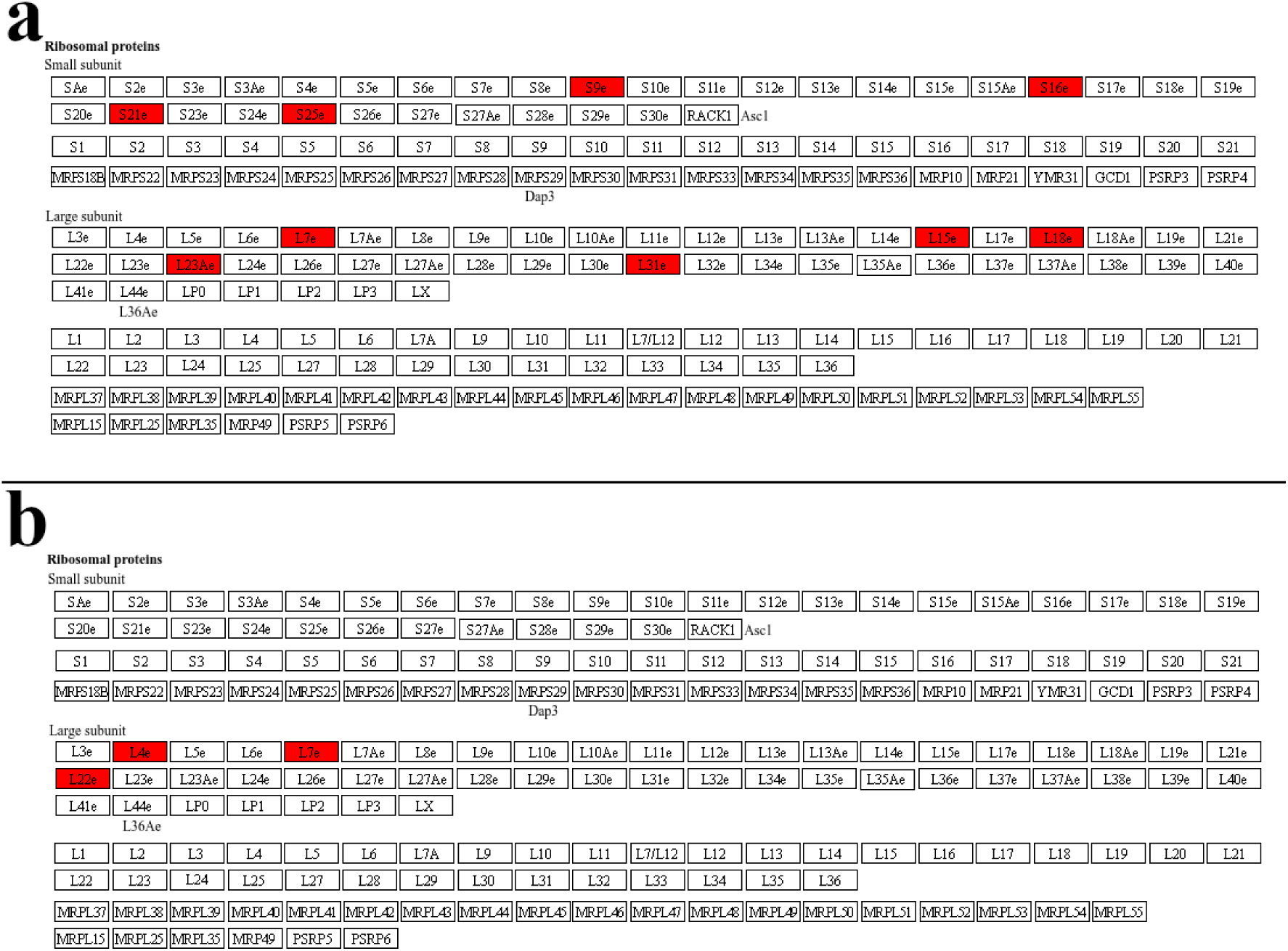
Comparison of ribosome KEGG pathway enrichment for the combined trait groups. Genes highlighted in red represent candidate genes identified from the ancestry-corrected GWAS. The figure displays a standardized KEGG reference map for the ribosome, which includes examples from different domains of life. (a) Ribosome pathway for the “yield & size” group. Numerous components of both the small (e.g., S9e, S16e) and large (e.g., L23Ae, L31e). (b) Ribosome pathway for the “color & pigmentation” group. A smaller, more specific set of ribosomal proteins (e.g., L4e, L7e, L22e) is highlighted, suggesting that pigmentation may be linked to specific regulatory functions of the ribosome.

### Corrected GWAS links distinct morphological traits to specific biological pathways

To demonstrate the breadth of biological mechanisms identified by our corrected GWAS, we highlighted the GO enrichment results for three distinct morphological traits (Fig. 7). The analysis for fruit shape - elliptic revealed an overwhelming enrichment for pathways related to nucleotide and ATP transport across mitochondrial membranes (Fig. 7a). This specific finding suggests that the developmental program creating an elliptic shape is strongly linked to the efficient transport and delivery of cellular energy (ATP) and genetic building blocks (nucleotides) to support localized growth.

**Fig. 7.**
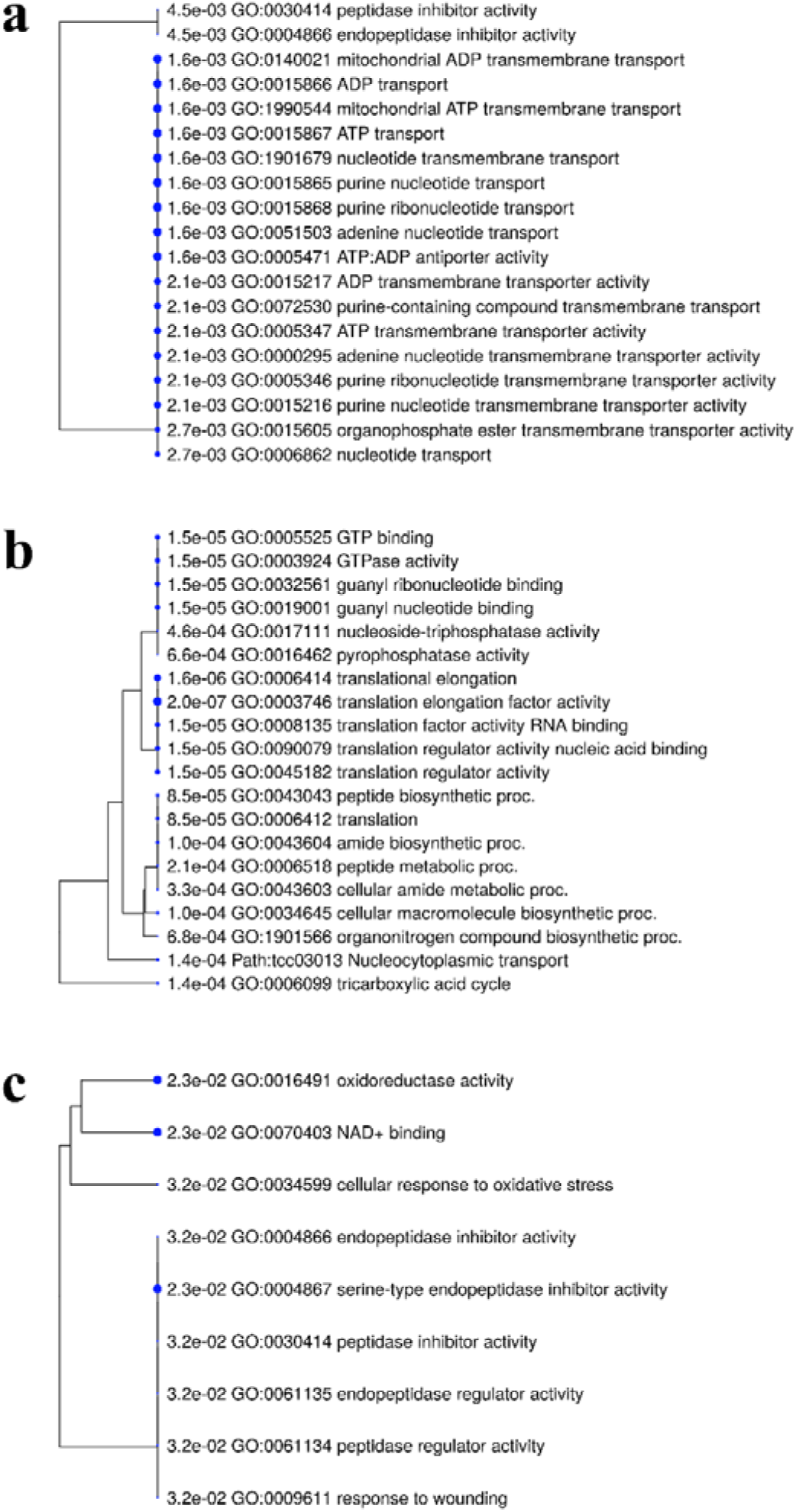
GO enrichment trees for notable individual morphological traits from the corrected GWAS. Each panel displays the top significantly enriched GO terms for a specific trait. (a) GO enrichment for fruit shape - elliptic, dominated by pathways for mitochondrial ATP and nucleotide transmembrane transport. (b) GO enrichment for fruit husk hardness, showing a strong association with the TCA cycle and the machinery of translation and protein synthesis. (c) GO enrichment for fruit apex form - attenuate, highlighting pathways related to cellular stress response and the regulation of protein activity.

In contrast, the analysis for Fruit Husk Hardness pointed to the fundamental machinery of protein synthesis and cellular energy production (Fig. 7b). The most significant terms were related to the tricarboxylic acid (TCA) cycle, which is the central hub of cellular respiration, and numerous terms related to translation and peptide biosynthesis.

Finally, the analysis for fruit apex form (‘attenuate’, a shape where the fruit’s tip gradually tapers to a long, slender point) implicated pathways involved in stress response and regulation (Fig. 7c). Enriched terms included cellular response to oxidative stress, response to wounding, and various peptidase inhibitor activities. This suggests that the development of this specific shape is tied to the genetic pathways that manage cellular stress and regulate protein turnover.

### Predictive modeling of wet bean mass using a Neural Network model

The pod index is calculated using the following formula:

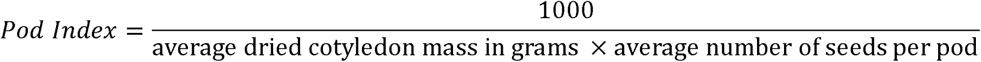

This calculation determines the number of pods required to produce one kilogram (1000 grams) of dried cocoa beans. A lower pod index value indicates a higher yield potential, as fewer pods are needed to produce the same amount of cocoa. Wet bean mass provides a more direct assessment of yield, so we employed a machine learning approach using Neural Network model to identify predictive phenotypic traits while excluding obvious correlating traits such as seed number, fruit length, fruit width, and fruit length to width ratio. The performance of the trained Neural Boosted model is shown in Fig. 8. The model exhibited strong predictive accuracy on both the training set (R-squared = 0.790 +/− 0.08) with and the validation set (R-squared = 0.715 +/− 0.12). The scatter plots of actual vs. predicted wet bean mass values show a close alignment of the points with the diagonal line representing perfect prediction, indicating that the model accurately captures the relationship between the predictor variables and wet bean mass. Further analysis using the Neural Boosted model identified cotyledon mass as the primary driver of wet bean mass, with a remarkably high importance score of 6 (Table S1). Cotyledon length emerged as the second most influential predictor, receiving an importance score of 2. Cotyledon length-to-width ratio showed slight predictive power with an importance score of 1, whereas cotyledon width and other phenotypic traits showed little to no predictive influence (Table S1). Cotyledon mass displayed a substantial positive influence on wet bean mass, evidenced by its main effect of 0.484 and total effect of 0.5198. This suggests that cotyledon mass impacts wet bean mass both directly and indirectly through interactions with other variables. Cotyledon mass (portion = 0.520) and cotyledon length (portion = 0.213) accounted for the vast majority (73.3%) of the predictive power in the model, highlighting their dominant roles in determining wet bean mass.

**Fig. 8.**
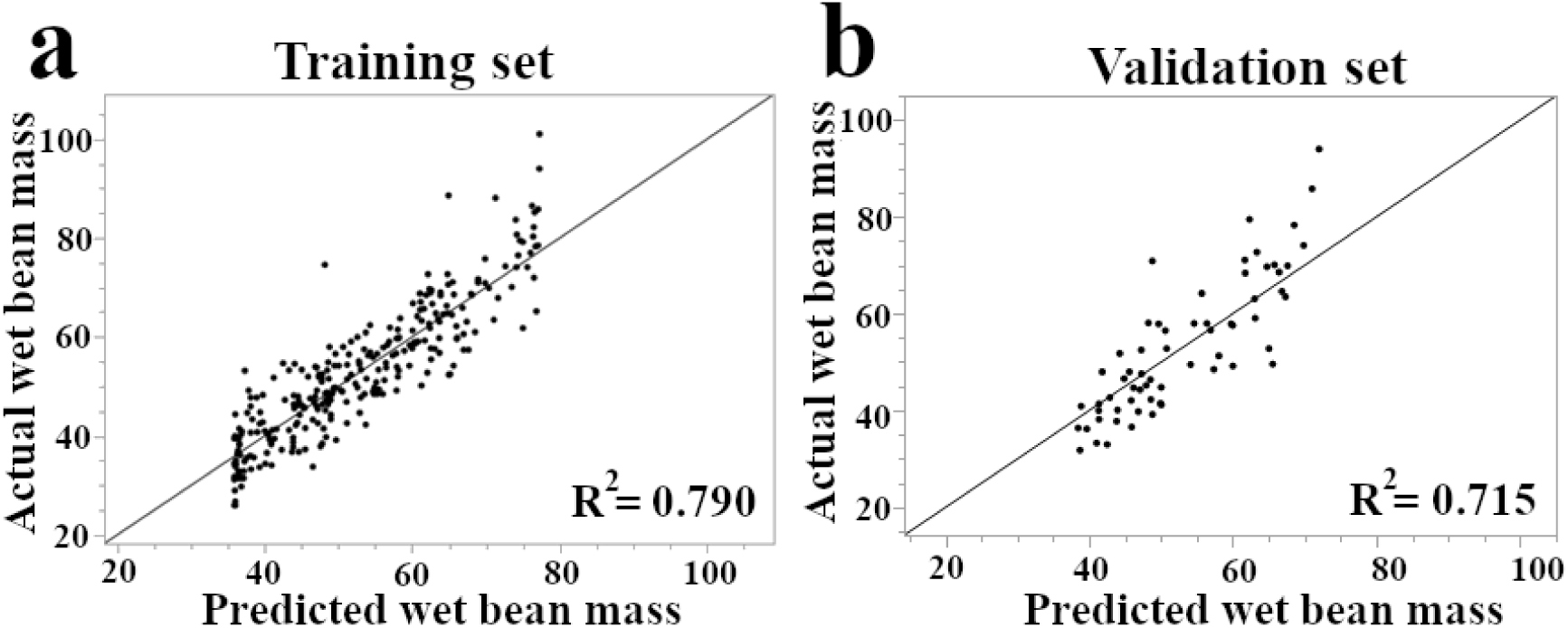
Evaluation of the Neural Boosted model for predicting wet bean mass in cacao, based on phenotypic data. The model was evaluated on the full dataset (n=346) using a robust 5-fold cross-validation procedure, which was repeated 20 times. (a) Actual vs. predicted values aggregated from the training folds. (b) Actual vs. predicted values aggregated from the validation folds. The black line in each plot represents a perfect prediction (actual = predicted). The R-squared values shown are the average performance across all validation runs, demonstrating the model’s generalized predictive accuracy.

## Discussion

### Correlation between patristic distances and phenotypic traits

We first examined how ancestry affected phenotypic traits in 344 diverse cacao accessions. The results indicate that broad ancestry accounts for a limited fraction of phenotypic divergence in this panel: significant traits show small positive patristic–phenotypic correlations, with the strongest for pod index (*r*= 0.25, q= 0.0019), followed by cotyledon width (*r*= 0.20, q= 0.0019). While genetic divergence has increased trait difference for these two traits, the weak overall correlation between patristic distances and phenotypic traits suggests that broad phylogenetic relationships are not the primary determinants of cacao phenotypic variation in this collection.

### Correcting for population structure reveals the core genetic basis of yield

A key methodological finding of this study was the impact of correcting for population structure in the GWAS, best illustrated by comparing the naive (uncorrected) and structured (corrected) models for the pod index trait. The naive analysis highlighted a top SNP in a peroxidase gene, a plausible but likely confounding signal for general plant vigor^23–25^ that could lead to spurious associations if population structure is not controlled^26,27^. In contrast, the corrected model elevated TcSNP597 to the top-ranked position, a locus within a gene for a 21 kDa proteinase inhibitor seed protein, a class of proteins known to be major determinants of seed quality and nutritional value^28^. This shift is significant, as it moves the focus from a general stress-response gene to a gene class with a specific role in seed development and defense, making it a more direct and plausible candidate. Similarly, the correction refined the associated pathway: the uncorrected results were enriched for the broad but relevant pathway of “protein processing in the endoplasmic reticulum,” while the corrected results pinpointed the core of this process, becoming overwhelmingly dominated by the “ribosome” pathway itself (Fig. 6).Together, these comparisons illustrate the utility of a dual-model approach, where the exploratory findings of a naive analysis are refined by a structured model to pinpoint the specific biological machinery, in this case, protein synthesis directly linked to the trait.

### The genetic architecture of cacao is centered on pleiotropic hubs for protein synthesis

Building on these robust results, our corrected GWAS revealed a core genetic network associated with cacao productivity, showing that a small set of loci are central to multiple yield-component traits. This phenomenon, known as pleiotropy, is a key feature of complex traits in major crops, where single loci often influence multiple, correlated yield components^29,30^. Specifically, TcSNP281, TcSNP597, and TcSNP1366 were all identified as top associations for combinations of pod index, wet bean mass, and seed number (Table 1). The putative genes at these loci are involved in fundamental biological processes: protein synthesis (60S ribosomal protein L31), seed development and defense (21 kDa proteinase inhibitor seed protein, known as a negative regulator for seed quality), and energy metabolism (heparan-alpha-glucosaminide N-acetyltransferase). The convergence of these crucial yield traits on the same genetic loci strongly suggests that they are regulated by a central hub controlling the plant’s capacity for protein production and energy allocation for seed development. This suggests that this network represents a key target of selection, where breeders have, either intentionally or unintentionally, selected for alleles that optimize the overall capacity for protein production to enhance yield. This finding aligns with a growing body of evidence highlighting that the synthesis of storage proteins and the regulation of amino acid and energy metabolism are tightly controlled, central processes that determine final seed composition and yield^31–34^.

The validity of our analytical approach is demonstrated by an internal positive control. The analysis for fruit surface anthocyanin recovered 6-phosphogluconate dehydrogenase (TcSNP174), a key enzyme biochemically upstream of flavonoid precursors. This finding, combined with the identification of a locus (TcSNP401) that converges with an independently reported finding for cacao pigmentation, provides strong support that our structured model successfully identifies pathway-specific, non-vigor associations. 6-phosphogluconate dehydrogenase catalyzes a rate-limiting step in the oxidative pentose phosphate pathway (OPP)^35^. The OPP pathway is critical as it generates carbon precursors for the shikimate pathway, a direct link that has recently been genetically demonstrated through the activity of 6-phosphogluconate dehydrogenase^36^. The shikimate pathway, in turn, is the well-established route for the biosynthesis of aromatic amino acids and subsequently the entire class of flavonoids, including the anthocyanin pigments that determine color^37,38^. Our ability to link a top statistical hit from our corrected model directly to a key, rate-limiting enzyme at the start of the relevant biochemical network provides strong validation for our findings.

### From predictive models to biological mechanisms

Our findings can be contextualized by comparing them to the initial analysis of this germplasm by Bekele et al.^11^. It is noteworthy that both studies, despite using different corrected GWAS models (Bootstrap Forest vs. MLM), independently identified TcSNP401 as a key locus associated with fruit surface anthocyanin. This independent validation strongly supports the importance of this genomic region in cacao pigmentation. Beyond this convergence, our machine learning-based approach identified a largely distinct set of top candidate genes for most traits, suggesting our analysis has captured different facets of the genetic architecture and provides a novel and complementary view of cacao yield genetics.

Furthermore, this study successfully created a bridge between predictive modeling and genetic markers. Our Neural Network model independently identified cotyledon mass and length as the most powerful predictors of final wet bean mass. Our GWAS then provided a genetic basis for these predictive traits. For instance, cotyledon length was strongly associated with a SNP in a gene for a mitochondrial cytochrome c oxidase subunit; this link is highly significant, as the activity of cytochrome c oxidase is known to be indispensable for proper plant embryogenesis and seed viability^39,40^. Cotyledon mass was associated with a SNP in a RING/U-box superfamily protein. This finding is well-supported by studies in other crops showing that this class of E3 ubiquitin ligases directly regulates seed and organ size by targeting key proteins for degradation^41,42^.

Our combined-trait GO analysis revealed the fundamental processes underpinning major phenotypic groups. By combining all nine yield- and size-related traits, the analysis converged on a single, overwhelmingly significant biological theme: the ribosome and protein synthesis. A deeper look revealed that numerous components of both the small and large ribosomal subunits were associated with these traits, suggesting that overall yield potential is linked to the general capacity and throughput of the entire protein synthesis machinery. This conclusion is well-supported by studies showing that the disruption of individual ribosomal protein genes leads to significant defects in plant growth, development, and seed size^43–45^. In parallel, the combined analysis of five color- and pigmentation-related traits pointed directly to primary energy metabolism, including glycolysis and ATP generation. A key pathway identified in this group was “photosynthesis - antenna proteins,” highlighting the specific gene *Lhcb4*. Antenna proteins regulate how much light energy reaches photosystem II and variations in these proteins can directly impact chloroplast redox balance which is a primary trigger for anthocyanins as photoprotective agents. This provides a direct mechanistic link between the machinery of light capture and the energetically expensive process of pigment production^46,47^.

Beyond these broad themes, the analysis of individual corrected traits revealed a diversity of specific biological mechanisms controlling morphology. For instance, the genetics of fruit shape - elliptic were not linked to general growth but to the specific machinery of ATP and nucleotide transport across mitochondrial membranes. This suggests a key role for “energy logistics” in developmental patterning, a concept supported by findings that disrupting mitochondrial transporters can severely hamper organ development and morphology^48,49^. In another example, fruit husk hardness was strongly associated with the Citrate (TCA) cycle and the machinery of translation. This provides a direct link between a physical property and the cell’s core metabolic and biosynthetic capacity, as studies have shown that restricting TCA cycle activity directly impairs the synthesis of secondary cell wall components like cellulose and lignin that confer structural rigidity^50,51^. These findings showcase the complexity of developmental genetics and demonstrate our approach’s ability to uncover novel and highly specific hypotheses for a wide range of traits.

Hierarchical clustering of traits based on the ancestry-adjusted GWAS results provided a global view of the genetic architecture, revealing several functionally significant groupings (Fig. 3). The analysis partitioned the traits into distinct clusters, most notably grouping key productivity metrics such as pod index, wet bean mass, seed number, and cotyledon length. This supports the concept of a shared genetic foundation for overall yield and corroborates the pleiotropic effects suggested by the single-trait GWAS, where the same SNPs were associated with multiple of these traits. Clustering of trait–gene association profiles underscored the modular nature of yield components. Key productivity metrics, pod index and wet bean mass, formed a tight pair and clustered with seed number (and ovule number, cotyledon length), consistent with a shared genetic backbone for overall yield. By contrast, cotyledon mass lay on a distant branch of the dendrogram, proximate to pod wall hardness, fruit length, and several apex□form traits. This separation indicates that, although phenotypically related, the genetic factors governing individual bean mass are partly distinct from those shaping bean count and pod□level characteristics. Practically, this suggests that bean size and bean number can be improved as semi□independent targets in selection schemes, rather than assuming one will automatically track the other.

Beyond the clustering of traits, our global analysis of SNP locations revealed a striking pattern in the underlying genetic architecture. By mapping the top three associated SNPs for all 27 traits, we identified several genomic “hotspots” containing associations for multiple, diverse traits (Fig. S2), a phenomenon that has been observed for important agronomic traits in other major crops^52,53^. A distinct trend emerged where many of these hotspots were concentrated in the subtelomeric regions of the chromosomes, particularly on Chromosomes 1, 2, 4 and 9. These subtelomeric regions are known to be dynamic parts of the genome, often characterized as being gene-rich and exhibiting high rates of recombination^54,55^. This unique combination of features has led to the hypothesis that subtelomeres act as “hotbeds” for new gene origination and are potential centers for rapid evolutionary adaptation^56,57^. This concentration of trait-associated loci suggests these dynamic genomic regions may have been key targets of selection during both the initial domestication of cacao and its subsequent, geographically diverse improvement.

### Implications for cacao breeding and future directions

The successful prediction of wet bean mass using a Neural Network model (Fig. 8), well-suited for capturing complex, non-linear relationships often found in biological systems, demonstrates the potential of leveraging phenotypic data for developing predictive tools for complex traits in cacao^58^. The model’s high accuracy (R-squared = 0.790 for training, 0.715 for validation) highlights the strong relationships between the predictor variables and wet bean mass. Notably, cotyledon mass and cotyledon length were identified as the two most important predictors (Table S1) of wet bean mass, a direct measure of yield. Larger cotyledons on young cacao seedlings may be indicators of vigor and higher photosynthetic capacity, which over the long-term could result in more productive cacao trees, a principle supported in other crops where early-stage growth traits have been shown to be significantly predictive of final yield^59^. While this suggests that simple measurement of these two traits could provide an accurate and cost-effective method for predicting productivity, further longitudinal evidence is required to demonstrate early screening utility of these key-traits. Nonetheless, the identification of these early-screening traits may have significant implications for cacao breeding programs that are often hampered by long generation times and an urgent need for new, high-yielding varieties^13^. Targeting specific combinations of traits, guided by the identified clusters (Fig. 3) and the machine learning model (Fig. 8), could lead to even more efficient selection strategies for improving yield potential. Furthermore, these results, combined with our GWAS findings, suggest that genomic selection, incorporating information from multiple loci across the genome, could be a powerful tool for accelerating cacao improvement, an approach particularly impactful for perennial crops with long breeding cycles^60^.

The findings of this study, particularly the successful application of a Neural Network model for predicting wet bean mass, provide a springboard for several promising avenues of future research in cacao. Given the potential impact on breeding strategies, prioritizing the validation of candidate genes identified through GWAS would be beneficial. This could involve gene expression studies and functional analyses using transgenic approaches or, more directly, CRISPR-Cas9-mediated gene editing, which is a critical step for validating the causal effect of candidate genes identified through association studies^61^. Second, expanding the association mapping and genomic selection studies to larger and more diverse populations of cacao, including different genetic groups and geographic origins, would help to confirm the identified associations and identify additional loci contributing to trait variation across a wider range of germplasm. This could also involve exploring genotype-by-environment interactions by evaluating the performance of diverse accessions in multiple locations in collaboration with international partners. Third, investigating the role of other types of genetic variants, such as copy number variations and structural variants, could improve predictive power. Fourth, integrating genomic data with other omics data, such as transcriptomic, proteomic, and metabolomic profiles, could further elucidate the molecular mechanisms underlying trait variation. Both the inclusion of structural variants and the use of multi-omics data are considered key future directions for enhancing genomic prediction models^60^. Moreover, exploring different Neural Network architectures, such as the deep learning methods that are increasingly being harnessed for genomic prediction, could provide even greater accuracy when modeling complex traits from high-dimensional data^62^.

This study provides a comprehensive analysis of the phenotypic and genotypic variation in a diverse cacao collection, dissecting the complex relationships between population structure, genetic markers, and key agronomic traits. A central outcome was the demonstration of the profound impact of population structure on association mapping. By applying an ancestry-adjusted model, our GWAS successfully filtered out misleading signals related to general plant ‘vigor’ to instead identify that the core genetic drivers of yield are robust, trait-specific pathways for ribosome and protein synthesis. This corrected approach led to the identification of a small set of loci which were responsible for influencing multiple yield components like pod index, wet bean mass, and seed number. The hierarchical clustering of these corrected results further illuminated the genetic landscape, partitioning traits into distinct functional groups related to yield and pigmentation. Moreover, a highly accurate machine-learning model identified cotyledon mass and length as the most powerful predictors of wet bean mass, a direct measure of yield. While this work delivers a simple and much needed early-stage screening tool for cacao breeding programs, further validation of cotyledon mass as a predictor of yield is required. Collectively, these findings enhance our understanding of the genetic architecture of yield in cacao and provide a powerful, statistically robust framework for developing superior varieties through modern breeding strategies.

## Materials and Methods

### Phenotype and genotype data

Phenotypic and genotypic data used in this study were obtained from a publicly available, large-scale assessment of cacao germplasm detailed in Bekele et al.^11^. The study utilized 421 cacao accessions from the ICGT, a diverse collection that includes 263 wild genotypes, 23 accession groups (e.g., wild cacao types such as AMAZ, GU, IMC, MO, NA, PA, POUND, RB, SCA and SPEC accession groups)^11,63^.

Phenotypic evaluations for 27 traits, encompassing flower, fruit, and seed characteristics, were derived from the dataset by Bekele et al.^11^. Flower traits included anthocyanin intensity in the column of the pedicel, sepal length, anthocyanin intensity on the ligule, ligule width, anthocyanin intensity in the filament, style length, and ovule number. Fruit traits included shape, basal constriction, apex form, surface texture, surface anthocyanin intensity in mature ridges, ridge disposition, primary ridge separation, pod wall hardness, fruit length, fruit width, and fruit length to width ratio. Seed traits included number, shape, cotyledon color, total wet bean mass, cotyledon length, cotyledon width, cotyledon mass, and cotyledon length to width ratio. Pod index, a key agronomic trait that reflects yield potential in cacao, was also included as a composite trait^11^.

Complete phenotypic data were available for 346 of the 421 accessions^11^. For the genotypic data, Bekele et al.^11^ targeted a total of 836 SNPs located in coding sequences, selected based on their similarity to known protein sequences. After excluding SNPs with low call rates, 671 high-quality SNPs were retained. Consequently, the final dataset used for analysis in this study consisted of 671 SNPs and phenotypic data for 346 accessions.

### Correlation between patristic distances and phenotypic traits

We tested whether more distantly related accessions are more phenotypically different by performing Mantel tests (Pearson, 9,999 permutations) between the patristic distance matrix and pairwise phenotypic distances for each trait. Geneious software (V 2025.2.1) was used to calculate patristic distances and GU305 and NA58 were omitted due missing overlaps resulting in a total of 344 out of 346 accessions. Natural log transformations were applied for normality on traits. FDR was controlled across traits.

### Analysis of population structure and phenotypic variation using PCA

To visualize the underlying structure of the phenotypic and genotypic variation among the cacao accessions, and to generate covariates for population structure correction, PCA was performed using JMP Pro 17. Prior to analysis, the categorical SNP data was processed. First, missing SNP genotypes (coded as ‘N’) were imputed using the Multivariate SVD Imputation method within the ‘Explore Missing Values’ platform of JMP Pro 17, using 6 singular vectors and a maximum of 10 iterations. Following imputation, the complete categorical SNP matrix was converted into a numerical format based on allele counts (e.g., 0, 1, 2) using JMP Pro’s built-in functionalities to serve as input for the analysis. Two separate PCAs were then conducted. The first was performed on the matrix of all measured phenotypic traits to visualize phenotypic relationships. The second was performed on the final imputed and numerically-coded SNP genotype matrix to assess population structure. From this genotypic PCA, the resulting Principal Component Scores for the first three components were saved back to the main data. To create the corrected phenotype for the GWAS, these three PCs were then used as predictors in a standard least squares linear regression model for each trait, and the resulting residuals were saved as the new, ancestry-corrected target variable.

### Machine learning-based GWAS to associate markers with traits

To identify genomic regions associated with phenotypic variation in the cacao accessions, we employed a machine learning GWAS approach based on Bootstrap Forest models implemented in JMP Pro 17. Unlike traditional linear models, which primarily test for additive effects of single markers, ensemble ML methods like Bootstrap Forest can capture complex, non-linear relationships and potential gene-gene interactions (epistasis), which is advantageous for dissecting polygenic traits.

To create a comprehensive view of the genetic architecture, two parallel analyses were conducted: a naive model without population structure covariates, and a structured association model that included the first three principal components as fixed-effect covariates. The naive model based analysis provides an exploratory view of the entire genetic landscape, identifying all associations including broad effects potentially confounded with population structure. In contrast, the structured association model based analysis statistically removes the ancestral background noise to pinpoint loci with a direct and independent effect on the trait. Comparing these two results allows for the identification of the most robust candidate genes and provides insight into how much of a trait’s genetic signal is tied to the underlying population structure.

The GWAS was conducted for all phenotypic traits, including those related to accession group, flower morphology, fruit morphology, seed characteristics, cotyledon and bean mass, and pod index. Pod index is a key agronomic trait that reflects yield potential, calculated as 1000/(average dried cotyledon mass × average number of seeds per pod)^11^. Importantly, it is a composite trait that takes into account both pod size and seed number, with lower pod index values indicating higher yield potential^11^. For the naive model based analysis, separate Bootstrap Forest models were trained to predict each raw phenotypic trait using the full set of SNP markers as predictors.

For the structured association model based, we first accounted for confounding due to population structure using a two-step method. A linear model was used to predict each trait using the first three principal components (PC1, PC2, and PC3) from the genotypic PCA as predictors. The model can be represented by the formula:

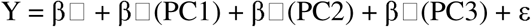

where Y is the vector of phenotypic values for a given trait, β□ is the overall mean, PC1, PC2, and PC3 are the vectors of the principal component scores, β□-β□ are their corresponding fixed-effect regression coefficients, and ε is the vector of residuals. The residuals from this model (ε), which represent the phenotypic variation independent of ancestry, were saved. Subsequently, separate Bootstrap Forest models were trained to predict these ancestry-corrected residuals for each trait using the full set of SNP markers as predictors.

For all Bootstrap Forest models in both analyses, default settings included the following parameters: number of trees in the forest was set to 100, the number of terms sampled per split was 1, the bootstrap sample rate was 1, the minimum number of splits per tree was 10, the maximum number of splits per tree was 2,000, and the minimum size split was 5. The top-ranking markers from both the corrected and uncorrected analyses were identified based on their importance scores, calculated as the “Portion” in JMP Pro. Given the high mapping resolution, with each marker representing a genomic region of approximately 1000 bp, we proceeded to identify the most likely candidate genes. To explore these potential candidates, we identified all annotated genes located within the genomic block represented by each top-ranking marker using the Criollo v2 reference genome (GCF_000208745.1).

### Gene ontology enrichment analysis

A GO enrichment analysis was conducted to investigate the potential biological functions and pathways overrepresented among the candidate genes associated with key agronomic traits. To compare the effects of population structure correction, this analysis was performed on gene lists generated from both the naive (uncorrected) and structured association (corrected) models for a subset of important traits, including pod index, wet bean mass, and anthocyanin intensity on the ligule.

For each analysis, a list of unique gene identifiers was compiled from all markers with a Portion score greater than 0.005, as identified by the Bootstrap Forest models. In cases where a marker could not be mapped to the reference genome, it was omitted from this downstream analysis. The curated list of unique *Theobroma cacao* gene IDs was submitted to ShinyGO v0.82^64^ (available at http://bioinformatics.sdstate.edu/go/). The enrichment analysis was conducted using the *Theobroma cacao* Belizean Criollo B97-61 B2 genome database. Default settings were applied, including a FDR cutoff of 0.05 to determine statistical significance. Pathway size filters were set to a minimum of 2 and a maximum of 5,000 genes per GO term.

### Hierarchical clustering of trait association profiles

To visualize the global genetic relationships between traits, we performed hierarchical clustering analysis. This analysis used the Ward method based on the importance scores (portion) derived from the structured Bootstrap Forest models for all traits. This approach groups traits that exhibit similar genetic association profiles, suggesting shared underlying genetic pathways or pleiotropic effects.

### Development of machine learning model for predicting wet bean mass

To further investigate the complex relationships between the various measured traits and wet bean mass, a machine learning approach employing Neural Network models was utilized^65^. The primary goal was to train a predictive model for wet bean mass, a key agronomic trait in cacao, based on the suite of phenotypic characteristics used in this study. Model selection and training were performed using JMP Pro 17. An initial model screening step was conducted using the built-in model screening function to evaluate the performance of various algorithms and identify the most promising model type for predicting wet bean mass. The Neural Boosted model emerged as the best-performing model and was therefore selected for further analysis. The model was configured with a single hidden layer containing three neurons with a hyperbolic tangent (TanH) activation function (NTanH(3)) and built using 20 boosting iterations (NBoost(20)).

To robustly assess the model’s predictive power, a repeated 5-fold cross-validation scheme was implemented on the full dataset (n=346). This procedure was repeated 20 times to ensure a stable and reliable estimate of the model’s performance. The Neural Boosted model, with the specified NTanH(3) and NBoost(20) settings, was trained and validated during each run of the cross-validation. Obvious correlating phenotypic traits that would lead to spurious associations were excluded such as seed number, fruit length, fruit width, and fruit length to width ratio, as to identify non-obvious phenotypic predictors. The performance of the trained model was assessed using the coefficient of determination (R-squared), averaged across all validation folds from the repeated runs. The importance of individual features (traits) in the Neural Boosted model was assessed to gain insights into the factors most strongly influencing wet bean mass prediction.

## Acknowledgements

We are also grateful to the reviewers for their constructive feedback. Mention of any trade names or commercial products in this article is solely for the purpose of providing specific information and does not imply recommendation or endorsement by the U. S. Department of Agriculture. USDA is an equal opportunity provider and employer, and all agency services are available without discrimination.

## Funding

This work is supported by the U.S. Department of Agriculture, Agricultural Research Service, In-House Projects No. 8042-21220-258-000-D and 8042-21000-303-000-D.

## Conflicts of interest

The authors declare no conflicts of interest.

## Author Contributions

E.A. conceptualized and designed the study. The investigation was performed by J.B., S.P., and E.A. Data analysis, validation, and visualization were conducted by J.B., D.L., S.P.C., S.P., and E.A. The methodology and resources were provided with contributions from J.B., I.B., S.L., J.H.J., S.P.C., A.H.L., and L.W.M. E.A. wrote the original manuscript draft. All authors reviewed, edited, and approved the final manuscript.

## Data availability

All phenotypic and genotypic data analyzed in this study are publicly available. These data were originally generated and published by Bekele et al. (2022) and can be found within the Supporting Information files associated with that publication (PLoS One 17(1): e0260907), accessible via: https://doi.org/10.1371/journal.pone.0260907.

## Supplementary Figures

**Fig. S1.**
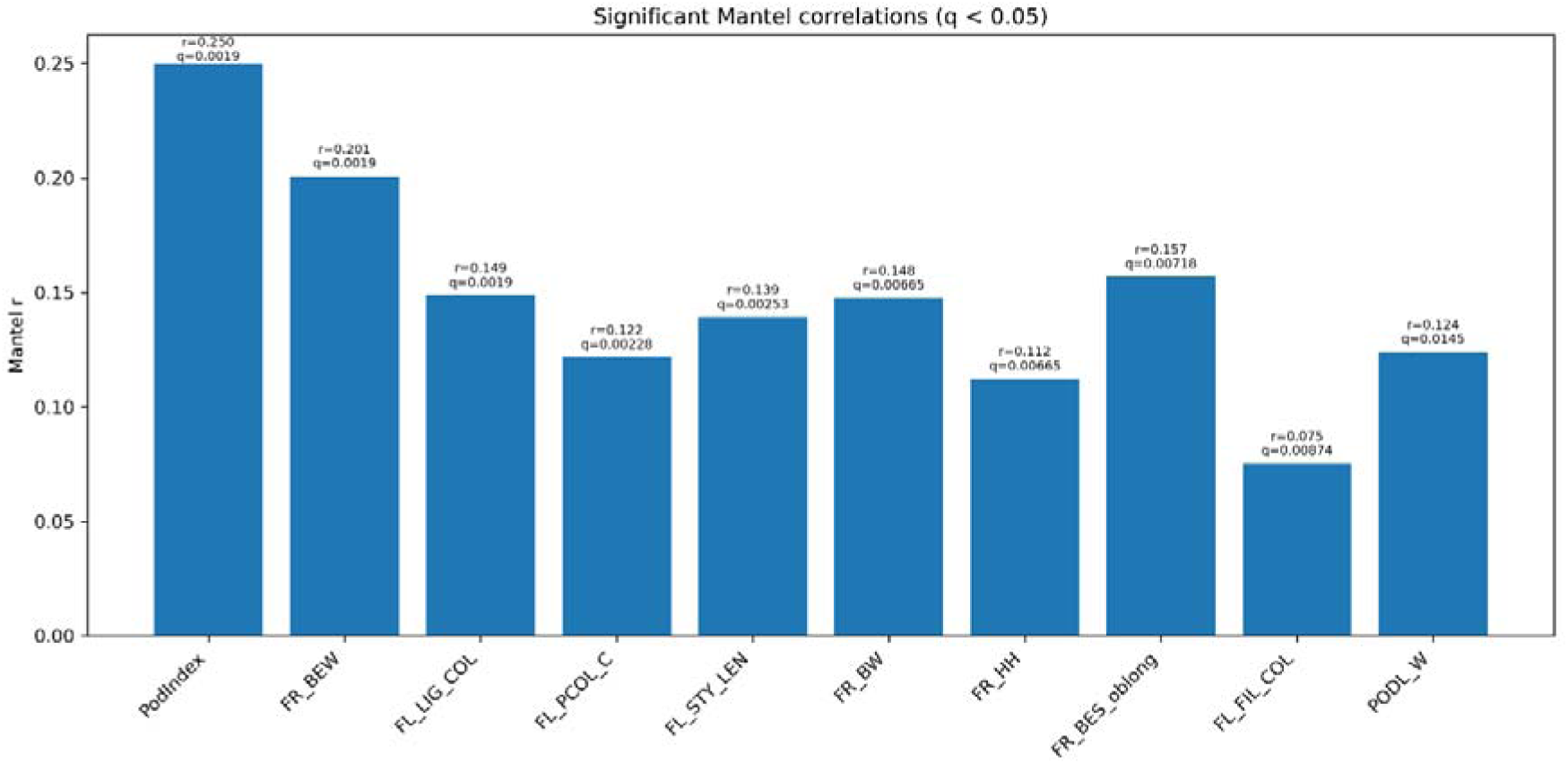
Bars show the Mantel correlation coefficient (r) between genetic distance and pairwise phenotypic difference for each trait. Only traits with q < 0.05 are displayed; bars are ordered by q and then r. Text above each bar reports r and q. Positive r indicates that more distantly related accessions tend to differ more in the trait. The categorical label “Acc Group” was excluded a priori. FR_BEW is cotyledon width, FL_LIG_COL is flower ligule anthocyanin intensity, FL_PCOL_C is flower pedicel column anthocyanin intensity, FL_STY_LEN is flower sepal length, FR_BW is cotyledon mass, FR_HH is pod wall hardness, FR_BES_oblong is seed shape oblong, FL_FIL_COL is flower style anthocyanin intensity, and PODL_W is pod length to width ratio.

**Fig. S2.**
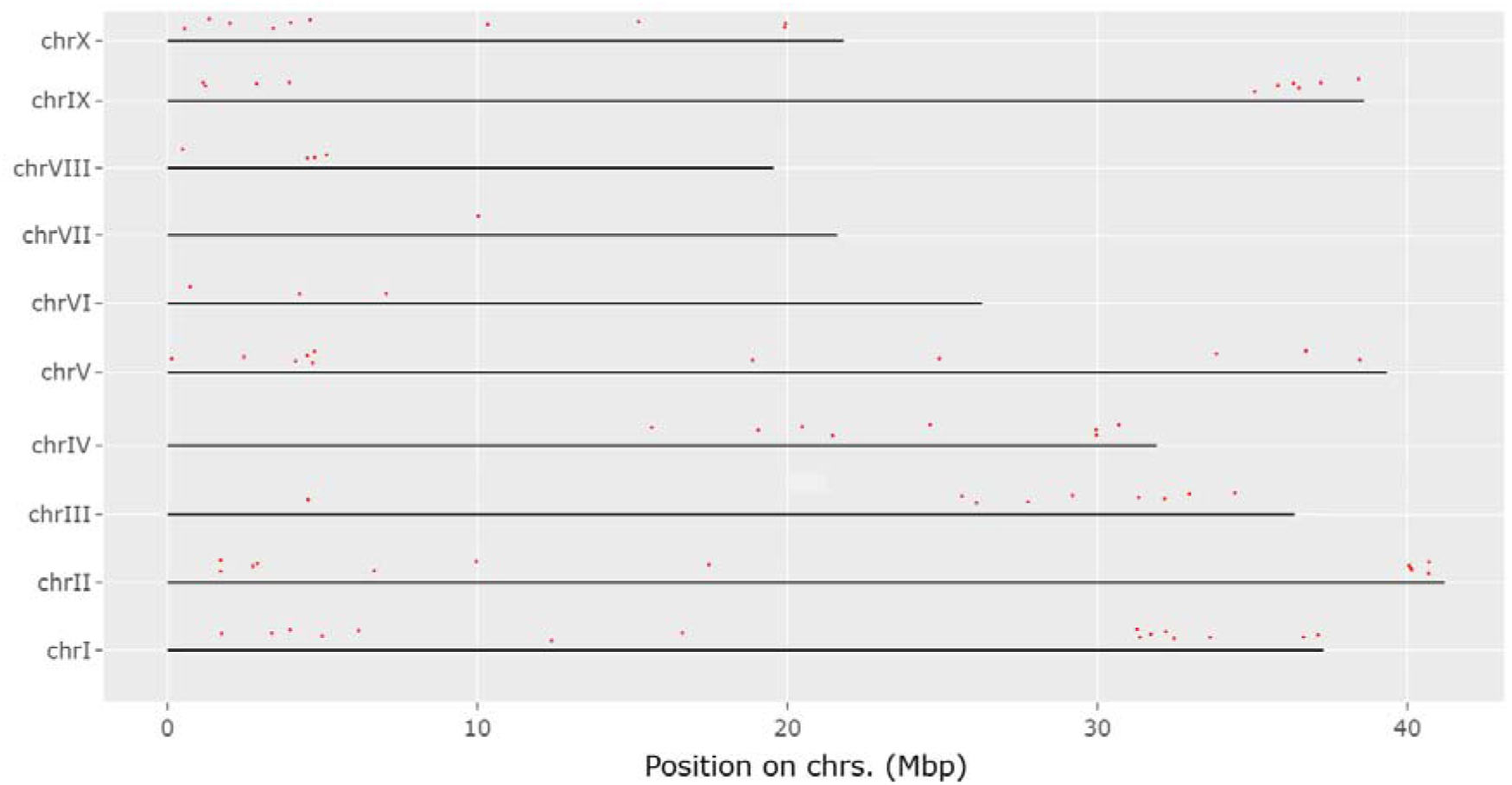
Genomic hotspot map of top associations across all traits. The plot shows the physical locations (Mbp) of the top 3 most significant SNPs for each of the 27 traits analyzed. Clear clusters, or “hotspots,” of associated loci are visible on several chromosomes. A complete list of all SNPs shown, their associated traits, and putative candidate genes can be found in Supplementary Data 1.

**Fig. S3.**
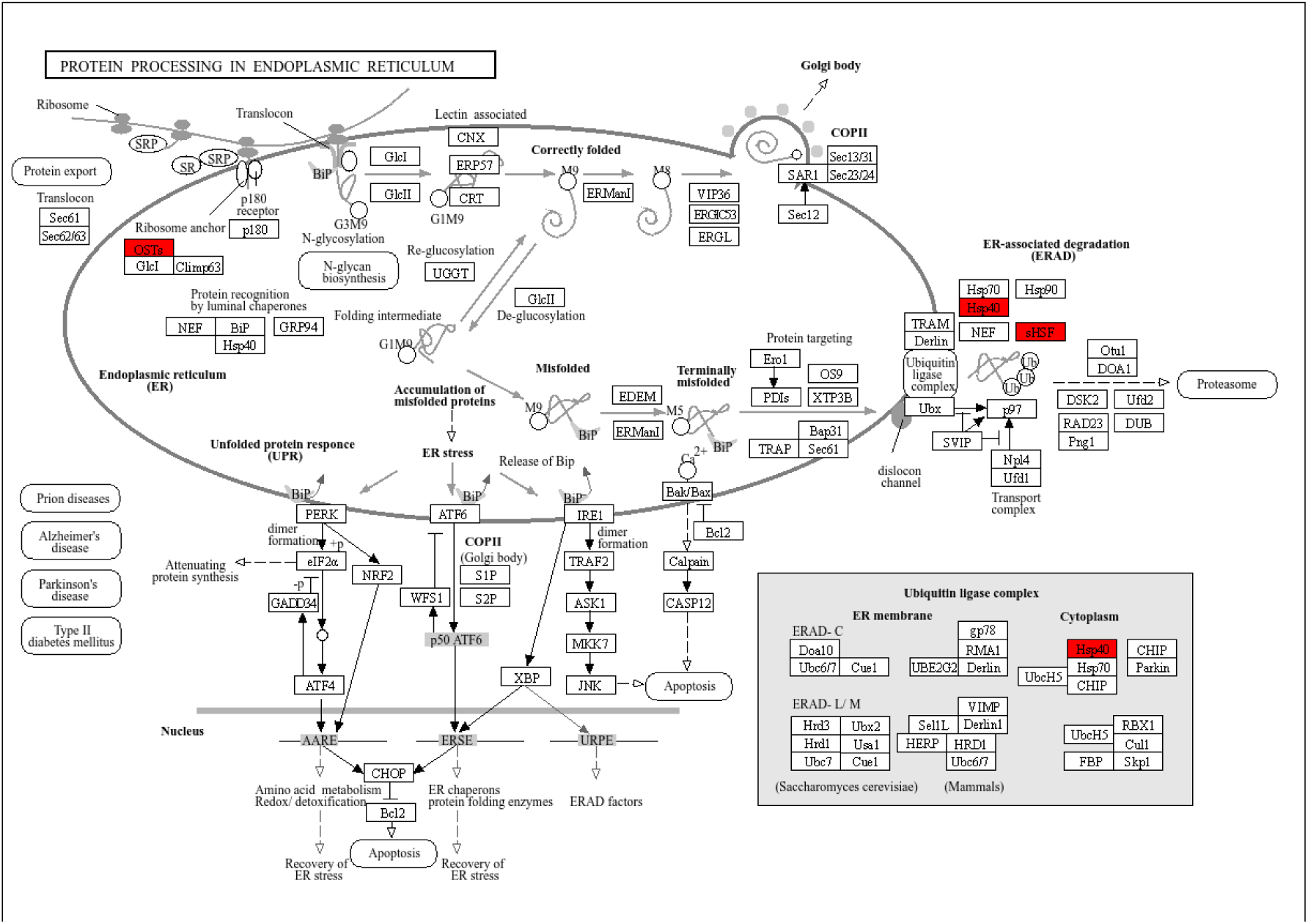
KEGG pathway for protein processing in the endoplasmic reticulum. This pathway was significantly enriched in the uncorrected GWAS for the pod index trait. The analysis identified several key genes within this pathway (highlighted in red), including components of the N-glycosylation machinery (*OST4*) and molecular chaperones involved in protein folding and the ER-associated degradation (ERAD) pathway (*Hsp90*, *Hsp40*). This suggests that protein quality control is a significant component of the broad “vigor” signal detected in the naive analysis.

**FIG. S4.**
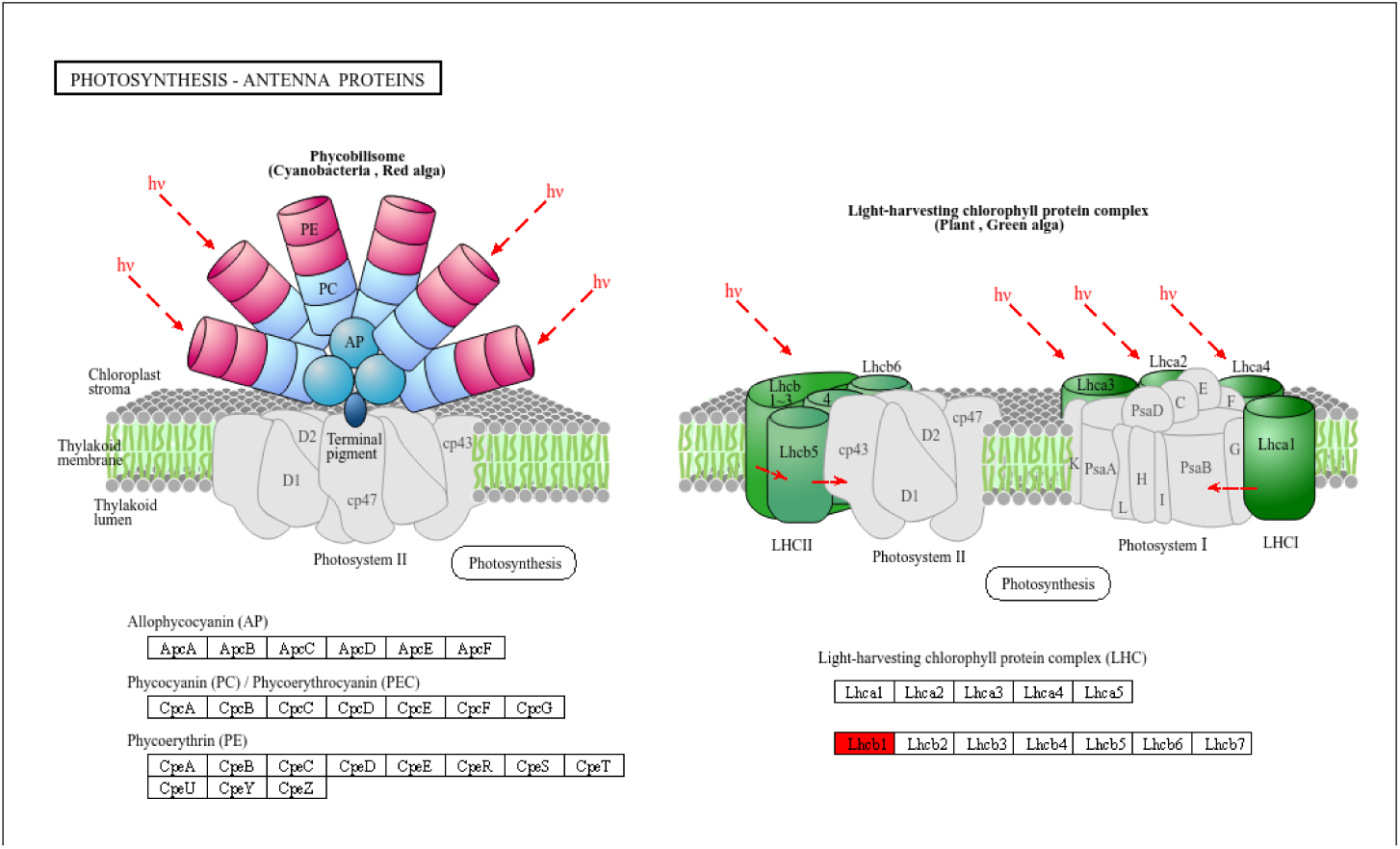
KEGG pathway for photosynthesis - antenna proteins. This pathway was identified as significantly enriched in the analysis of the pod index trait for both the naive (uncorrected) model and the structured association (corrected) model, which highlighted a key gene, *Lhcb1* (highlighted in red), encoding a major light-harvesting chlorophyll-binding protein of photosystem II. The consistent significance of this pathway across both analytical models suggests that the efficiency of light capture is a robust component of the genetic architecture for yield-related traits in this cacao collection.

**Fig. S5.**
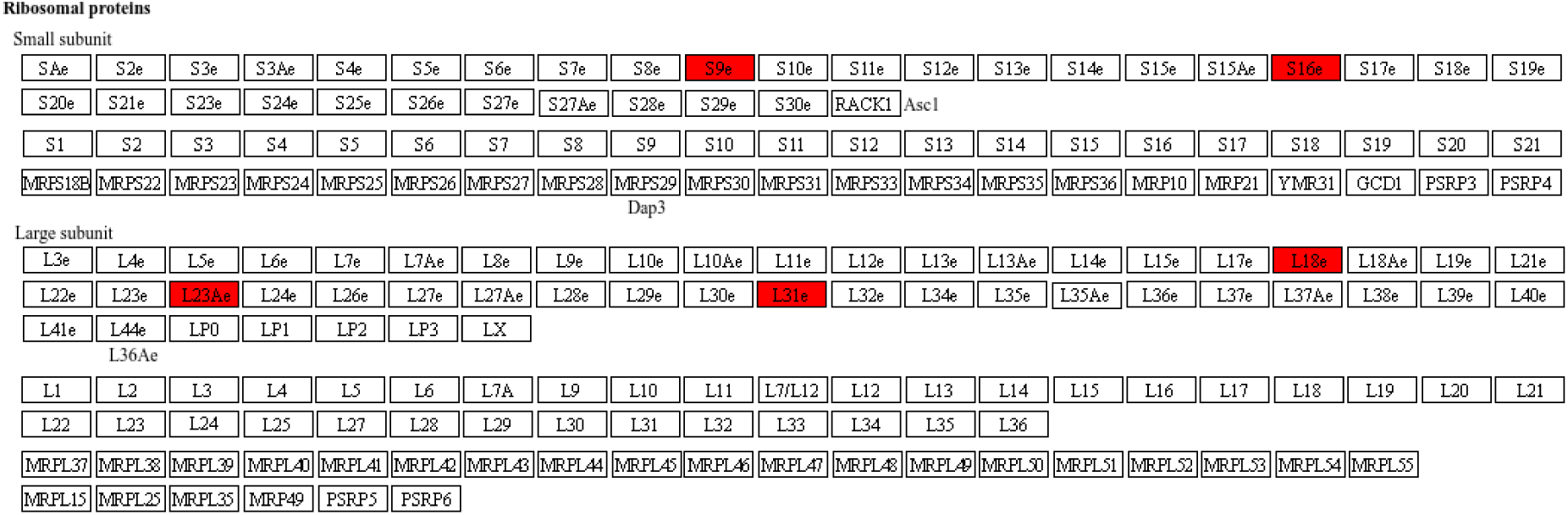
KEGG pathway for the ribosome. The pathway was the most significantly enriched result from the structured association (corrected) GWAS for the pod index trait. The analysis identified multiple genes encoding specific ribosomal proteins (highlighted in red), including components of the small subunit (*S9e*, *S16e*) and the large subunit (*L23Ae*, *L31e*, *L18e*). This finding strongly suggests that the genetic variation controlling yield potential, after correcting for population structure, is directly linked to the core machinery of protein synthesis.

**Fig. S6.**
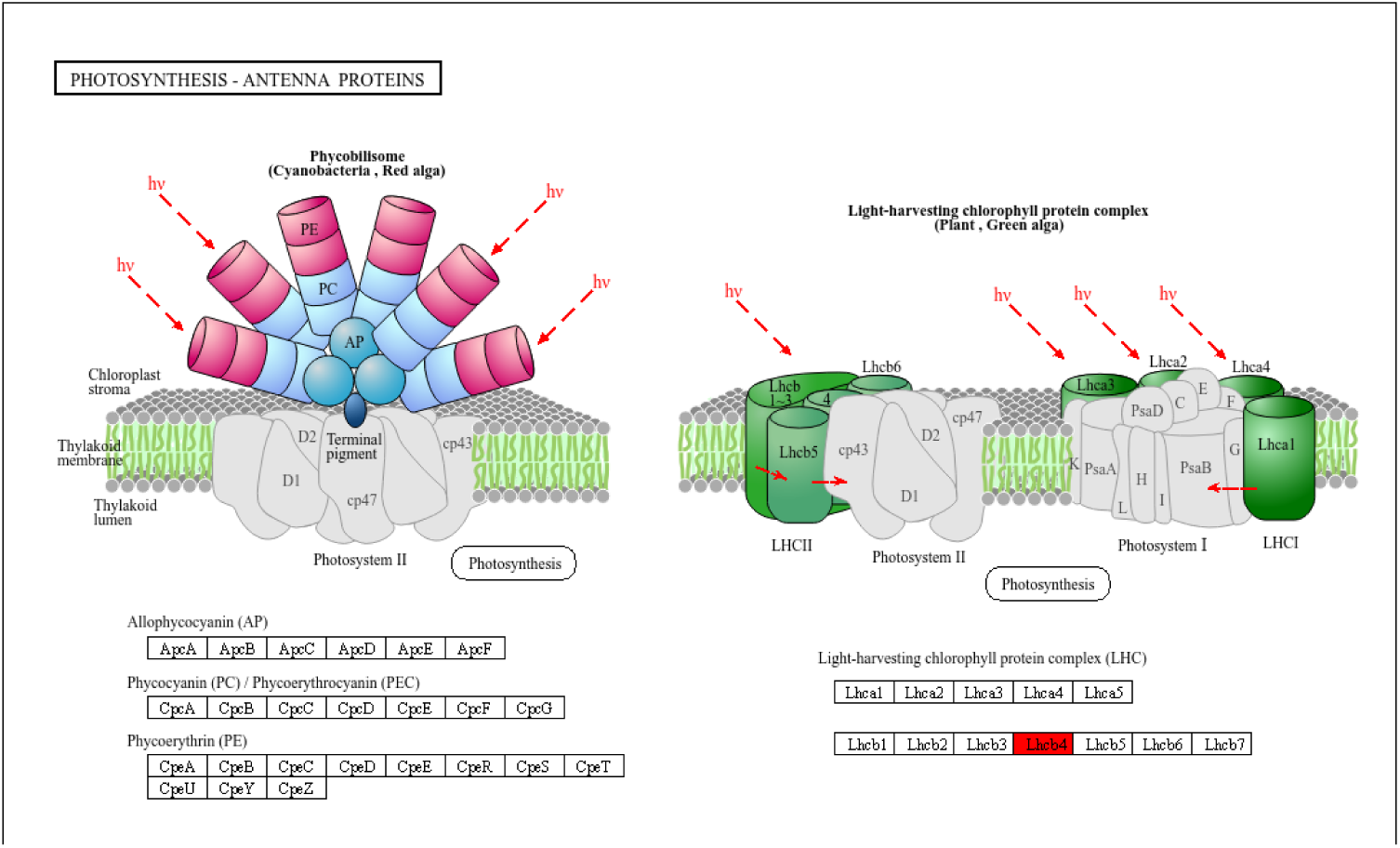
KEGG pathway for photosynthesis - antenna proteins. The pathway was identified as significantly enriched in the corrected, combined-trait analysis for the “color & pigmentation” group. The analysis highlighted the gene *Lhcb4* (highlighted in red), which encodes a key component of the light-harvesting chlorophyll protein complex. This finding suggests a specific link between the machinery of light capture and the energy-intensive process of producing the pigments responsible for color.

## Supplementary Tables

**Table S1.** Most important traits for wet bean mass prediction based on the Neural Boosted model analysis. Relative importance of the top four predictor traits as determined by the Neural Network model. Feature importance was assessed using a repeated 5-fold cross-validation procedure on the full dataset of 346 cacao accessions. The table lists each trait, its main effect, total effect, and importance score. The importance score reflects the contribution of each trait to the model’s predictive accuracy, with higher scores indicating a stronger influence. Cotyledon mass was identified as the most important predictive variable.

| Trait | Main effect | Total effect | Importance score |
| --- | --- | --- | --- |
| Cotyledon mass | 0.4843 | 0.5198 | 6 |
| Cotyledon length | 0.1864 | 0.2133 | 2 |
| Cotyledon length to width ratio | 0.0867 | 0.1101 | 1 |
| Cotyledon width | 0.04815 | 0.06895 | 0 |

## Notes

### Competing Interest Statement

The authors have declared no competing interest.

### Summary of Updates

Revised potential errors and conducted further analysis.

